# Fetal Kidney Transplantation for In Utero Fetuses

**DOI:** 10.1101/2024.04.15.589452

**Authors:** Keita Morimoto, Shuichiro Yamanaka, Kenji Matsui, Yoshitaka Kinoshita, Yuka Inage, Shutaro Yamamoto, Nagisa Koda, Naoto Matsumoto, Yatsumu Saito, Tsuyoshi Takamura, Toshinari Fujimoto, Shohei Fukunaga, Susumu Tajiri, Kei Matsumoto, Katsusuke Ozawa, Seiji Wada, Eiji Kobayashi, Takashi Yokoo

**Affiliations:** Division of Nephrology and Hypertension, Department of Internal Medicine, The Jikei University School of Medicine; Department of Urology, Graduate School of Medicine, The University of Tokyo; Department of Pediatrics, The Jikei University School of Medicine; Department of Urology, The Jikei University School of Medicine; Center for Maternal-Fetal, Neonatal and Reproductive Medicine, National Center for Child Health and Development; Department of Kidney Regenerative Medicine, The Jikei University School of Medicine

**Author notes:** Correspondence author: Takashi Yokoo. email.

## Abstract

Potter sequence, characterized by bilateral renal agenesis, oligohydramnios, and consequent pulmonary hypoplasia, presents a significant challenge in the management of affected neonates. Due to their prematurity and associated abdominal complications, these infants often fail to reach a stage where dialysis can be safely initiated and sustained, leading to an exceedingly high mortality rate. Therefore, there is hopeful anticipation that interventions serving as a bridge to achieve a state where dialysis can be safely performed will markedly improve life expectancy. We have developed a unique approach of “transplantation of fetal kidneys from a different species during the fetal period” as a bridge therapy until stable dialysis therapy can be implemented. This is a new concept of fetal therapy, targeting the fetus in utero and utilizing fetal kidneys of an appropriate size for transplantation.

In this study, we first validated the approach using allogeneic transplantation. Fetal kidneys with bladders from GFP-expressing rats (gestational age 14.0-16.5 days) were transplanted subcutaneously into allogeneic rat fetuses in utero (gestational age 18.0-18.5 days) using a special needle transuterinally, and live pups were successfully obtained. The transplanted fetal kidneys with bladders were confirmed to have urine production capability. By periodic aspiration of the subcutaneous urinary cyst after birth, urine produced by the transplanted fetal kidney was successfully drained outside the body for an extended period (up to 150 days). Biochemical tests confirmed the solute removal capacity of the transplanted fetal kidney. Furthermore, despite allogeneic transplantation, long-term urine production was sustained without the use of immunosuppressants, confirming that organ transplantation into fetuses is associated with lower rejection compared to adult transplantation. Next, xenotransplantation was performed. When GFP-expressing mouse fetal kidneys (gestational age 13.0-13.5 days) were transplanted into rat fetuses in utero, maturation of renal tissue structures was confirmed even in the interspecies setting.

## 1. Introduction

Potter sequence, characterized by bilateral renal agenesis, oligohydramnios, and consequent pulmonary hypoplasia, is one of the most serious perinatal diseases 1,2,3,4). Bilateral renal dysplasia, resulting in Potter sequence, occurs in 1 in 4000 individuals 5,6). In such cases, either fetal death occurs, or all affected infants die within 12 hours after birth due to pulmonary hypoplasia 6). Even if pulmonary hypoplasia is prevented by continuous amniotic fluid infusion therapy and immediate postnatal death is avoided, dialysis therapy is essential from birth. However, due to their prematurity and associated abdominal complications, these infants often fail to reach a stage where dialysis can be safely initiated and sustained, leading to an exceedingly high mortality rate 7). Therefore, there is hopeful anticipation that interventions serving as a bridge to achieve a state where dialysis can be safely performed will markedly improve life expectancy. Currently, there is no established effective treatment for such severe fetal renal failure, and dialysis therapy is performed as a limited treatment option after birth. We have previously reported that rat fetal kidneys can grow, differentiate, and produce urine in the body (retroperitoneal space) of adult rats 8,9,10,11,12). Similar phenomena have been confirmed in interspecies combinations such as rat-mouse and pig-mouse 13,14). We have developed the transplantation of the fetal kidney, ureter and bladder unit as a method to avoid hydronephrosis for a limited period 8). The urological unit was referred to as fetal kidneys with bladders (metanephroi with bladders: MNBs). Fetal kidneys at the stage of development that we are considering as a donor organ have not yet formed a glomerular structure with vascular invasion 15,16). Therefore, after transplantation, recipient-derived blood vessels invade the transplanted fetal kidneys and form glomeruli. Vascular endothelium is at the forefront of rejection in organ transplantation, and the advantage of constructing the vascular endothelium of the transplanted organ with self-vessels is that the vessels are less likely to be the target of rejection 17,18).

Therefore, we considered transplanting xenogeneic MNBs into intrauterine fetuses as a bridge until dialysis can be stably performed. However, it is unclear whether MNBs can be transplanted, engrafted, and mature in the fetus targeted for treatment, and there have been no reported cases. In this study, we transplanted rat MNBs into the subcutaneous tissue of rat fetuses and confirmed their engraftment, maturation, and urine production. We also investigated the immunological advantages of using fetuses at the developmental stage of the immune system as recipients. In addition, the model of xenogeneic transplantation was validated by transplanting mouse fetal kidneys into rat fetuses, confirming the maturation of the transplanted kidneys and demonstrating less tissue damage due to rejection compared to the transplantation of mouse fetal kidneys into adult rats. We aim to develop an innovative treatment for severe fetal renal failure and have conceived a completely new concept of fetal therapy: transplanting xenogeneic MNBs during the fetal period. This study is the world’s first report demonstrating the effectiveness of organ transplantation during the fetal period and is expected to contribute to the development of groundbreaking treatments for congenital kidney diseases, like Potter sequence. Although there have been reports of transplanting smaller cells into the amniotic fluid 19), peritoneal cavity 20,21), or retroperitoneal space 22) of intrauterine fetuses, to our knowledge, there have been no reports of transplanting larger tissue bodies, such as organs, into intrauterine fetuses and further endowing them with function, including in rodent models. This study is the first in the world to succeed in transplanting fetal organs during the fetal period and to analyze the transplanted tissue.

## 2. Results

### 2.1 Transplant GFP-SD rat MNBs into the SD rat fetal subcutaneous space

We developed a method for transplanting GFP (Green Fluorescent Protein)-SD (Sprague-Dawley) rat MNBs into the subcutaneous space of rat fetuses in utero. An overview of the experiment is shown in Fig.1a. One extracted GFP-SD rat MNB (E14.0-16.5) (Fig.1b) was loaded onto the tip of a 15-16G needle. Pregnant SD rats (E18.0-18.5) were laparotomized through a midline incision, and the uterus was gently exteriorized from the abdominal cavity. The position of the fetuses was confirmed through the semi-transparent uterine wall (Fig.1c). The uterine wall was punctured with the 15-16G needle loaded with GFP rat MNBs on the dorsal side of the fetus, and the needle was further inserted into the fetal subcutaneous space while being monitored under a stereomicroscope (Fig.1d,e). Approximately 0.1-0.5 mL was ejected to transplant the GFP rat MNBs loaded in the needle tip into the fetal subcutaneous space. Immediately after transplantation, the presence of GFP-positive tissue was confirmed using a fluorescence stereomicroscope to ensure proper transplantation (Fig.1f). To minimize the duration of surgery, 2-4 fetuses per mother were used as recipients (approximately 40 minutes of total anesthesia time for transplantation into 4 fetuses). The uterus was gently returned to the abdominal cavity, and the incision was closed. Four days after transplantation, on embryonic day 22, natural delivery occurred, and live offspring were successfully obtained. By puncturing the subcutaneous space of the fetuses in utero, the invasiveness was reduced, and the fetuses were able to survive the invasiveness of the thick 15-16G needle. The survival rate by natural delivery after transplantation surgery was 76%. On the day of natural delivery, the backs of all offspring were photographed using a fluorescence stereomicroscope to confirm the presence of GFP-positive tissue. Among the transplanted fetuses, GFP-positive tissue was confirmed in offspring with an average transplant success rate of 88% (Table.1).

**Figure 1.**
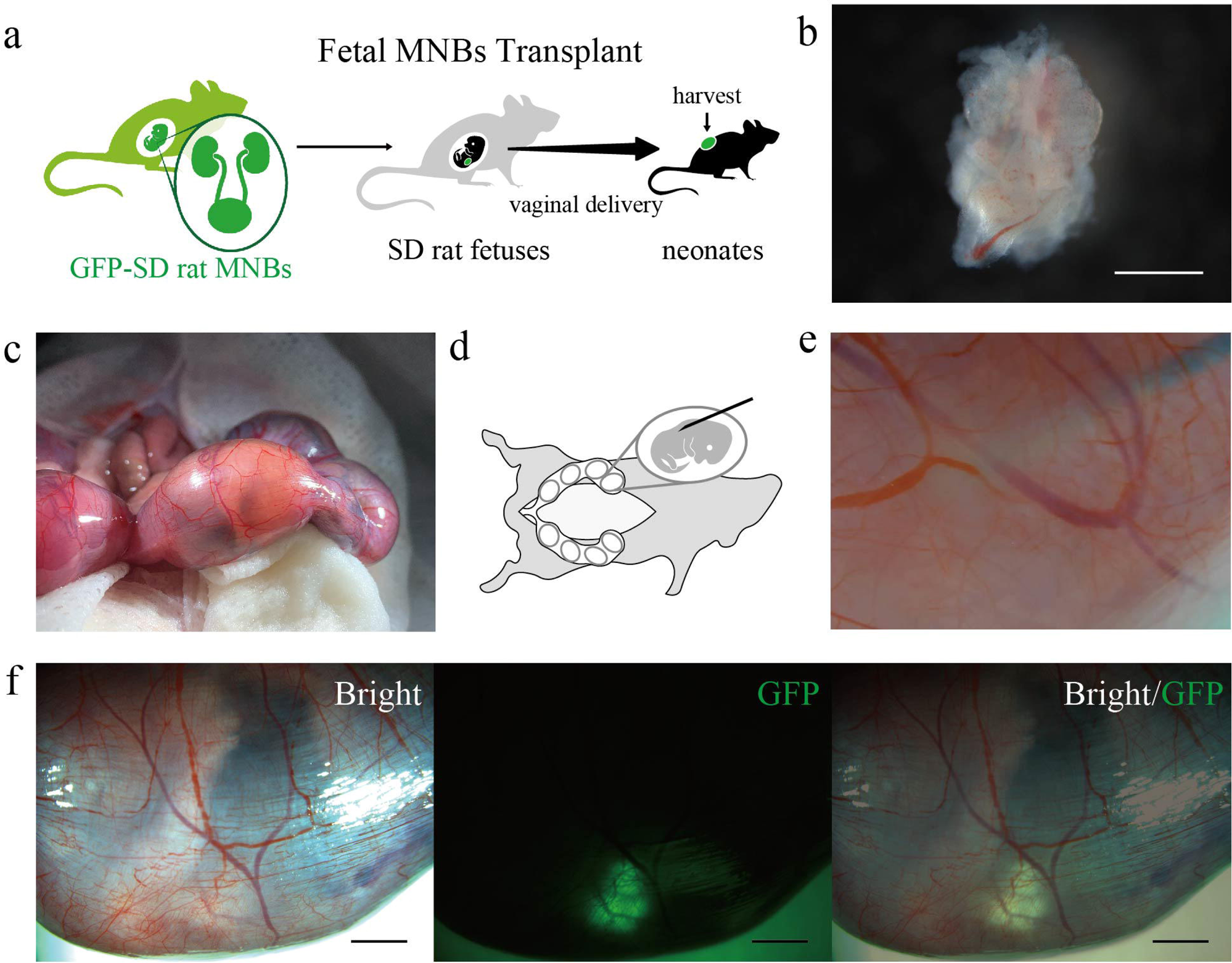
Transplantation method of MNBs into the fetus. a A summary of this experiment is shown. Transplanting GFP-SD rat MNBs (E14.0-16.5) into SD rat fetuses and harvesting MNBs after birth. b Representing the comprehensive profile of GFP-SD rat MNBs (E14.5). c The position of the fetuses was identified through the translucent uterine wall. d Depicting the image of exposing the uterus and puncturing the fetus. e The needle was inserted directly into the fetal subcutaneous space. f The presence of GFP-positive tissue was confirmed by fluorescence stereomicroscopy immediately after transplantation. Scale bars, 1 mm in (b); 2 mm in (f).

**Table 1.** Transplant success rate.

|  | **Number of**<br>**live pups/**<br>**All fetuses** | **Survival**<br>**rate[%]** | **Number of**<br>**fetuses**<br>**transplanted** | **Number of**<br>**GFP positive**<br>**neonates** | **Success**<br>**rate[%]** |
| --- | --- | --- | --- | --- | --- |
| Case1 | 12/14 | 86 | 2 | 1 | 50 |
| Case2 | 9/11 | 82 | 4 | 4 | 100 |
| Case3 | 11/14 | 79 | 2 | 2 | 100 |
| Case4 | 8/14 | 57 | 1 | 1 | 100 |
| Average | – | 76 | – | – | 88 |

### 2.2 Maturation of transplanted GFP-SD rat MNBs in neonates

After natural delivery on embryonic day 22, four days after GFP-SD rat MNBs transplantation, the neonates were nursed by their biological mothers and grew. There was no significant difference in body weight gain between the transplanted (n=3) and non-transplanted (n=3) neonates, indicating no growth impairment associated with the transplantation (Fig.2a). GFP-positive tissue was observed from the body surface of the neonates using a fluorescence stereomicroscope up to 28 days post-transplantation (24 days after birth), and a gradual increase in the size of the transplanted MNBs was noted (Fig.2b,c). On day 28 post-transplantation, the dorsal epidermis was incised, and the transplanted MNBs were examined, confirming the invasion of recipient subcutaneous blood vessels into the MNBs (Fig.2d). Percutaneous ultrasonography revealed the presence of a urinary cyst (Fig.2e), suggesting that the transplanted MNBs were producing urine. Histologically, glomeruli containing Nephrin-positive cells, LTL(Lotus tetragonolobus lectin)-positive proximal tubules, and ECAD(E-cadherin)-positive distal tubules were observed, confirming the maturation of the MNBs (Fig.2f,g). Additionally, GFP-negative CD31-positive vascular endothelial cells were found, suggesting that recipient-derived blood vessels invaded and formed glomeruli, producing urine (Fig.2h,i). On the other hand, GFP-positive CD31-positive blood vessels were also observed, indicating the presence of donor-derived vasculature (Fig.2j). These findings demonstrate the successful creation of a mature exogenous kidney by transplanting allogeneic MNBs into rat fetuses in utero.

**Figure 2.**
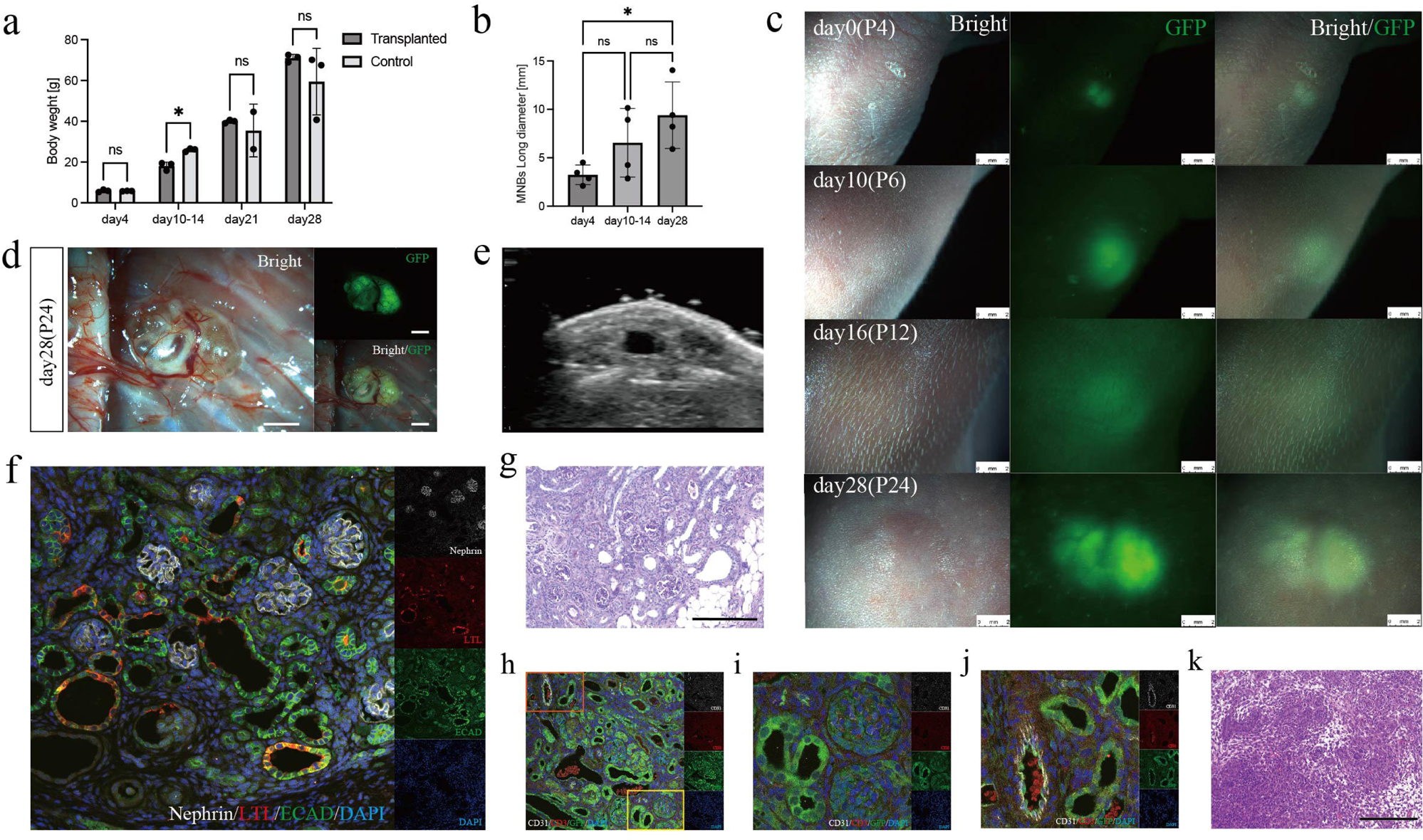
Maturation of transplanted GFP-SD rat MNBs in neonates. a After natural delivery on embryonic day 22, there was no significant difference in body weight gain between the transplanted (n=3) and non-transplanted (n=3). b,c GFP-positive tissue was observed from the body surface of the neonates and a gradual increase in the size of the transplanted MNBs was noted (n=4). All data are presented as mean ± standard error of the mean. The data were analyzed using the two-tailed unpaired t-test. *p<0.05. d On day 28 post-transplantation, recipient subcutaneous blood vessels had invaded the transplanted MNBs. e Percutaneous ultrasonography revealed the presence of a urinary cyst, suggesting that the transplanted MNBs were producing urine. f Histologically, glomeruli containing Nephrin-positive cells, LTL-positive proximal tubules, and ECAD-positive distal tubules were observed, confirming the maturation of the MNBs. g On day 28 post-transplantation, mature glomerular and tubular structures were confirmed by periodic acid-Schiff staining. h The yellow frame represents Fig.2i, while the orange frame represents Fig.2j. i The vessels within the glomerulus (CD31-positive) are GFP-negative, indicating recipient-derived vessels. J The vessels in the interstitial area (CD31-positive) are GFP-positive, indicating donor-derived vessels. k Hematoxylin and eosin staining. When GFP-SD rat MNBs were transplanted into adult SD rats and the tissue was retrieved 14 days post-transplantation, rejection was observed to the extent that glomerular and tubular structures could not be identified. Scale bars, 2 mm in (c) and (d); 200μm in (g) and (k). GFP, green fluorescent protein; LTL, Lotus tetragonolobus lectin; ECAD, E-cadherin.

Furthermore, when GFP-SD rat MNBs were transplanted into adult SD rats and the tissue was retrieved 14 days post-transplantation, rejection was observed to the extent that glomerular and tubular structures could not be identified (Fig.2k). Compared to transplantation into adults, MNBs transplantation into fetuses showed reduced rejection. This suggests that MNBs transplantation into fetuses may have immunological advantages regarding both GFP antigen and allogeneic grafts.

### 2.3 Development of the urine excretion method for GFP-SD rat MNBs transplanted into subcutaneous space

GFP-SD rat MNBs were transplanted into SD rat fetuses. Four days after transplantation, on embryonic day 22, natural delivery occurred. At 22 days post-transplantation (18 days after birth), a clearly raised structure, presumably the transplanted MNBs, was observed from the body surface (Fig.3a). Percutaneous ultrasonography revealed a distinct fluid accumulation with a long diameter of approximately 1 cm, indicating sufficient urine production capacity for safe percutaneous aspiration (Fig.3b). Subsequently, a single puncture was performed from the back using a 23-29G needle, and urine was aspirated (Fig.3c). Thereafter, single punctures were repeated 1-2 times per week, and continuous and stable urine excretion to the outside of the body was successfully maintained up to 150 days post-transplantation (146 days after birth) (Fig.3d). On average, it was shown that approximately 1 mL of urine was produced per day. Additionally, creatinine clearance (CCr) was measured, and it ranged from approximately 40 to 80 μL/min, confirming that the produced fluid was urine with solute removal capacity (Fig.3d). After confirming long-term urine excretion, tissue retrieval was performed at 150 days post-transplantation (146 days after birth) (Fig.3e,f).

**Figure 3.**
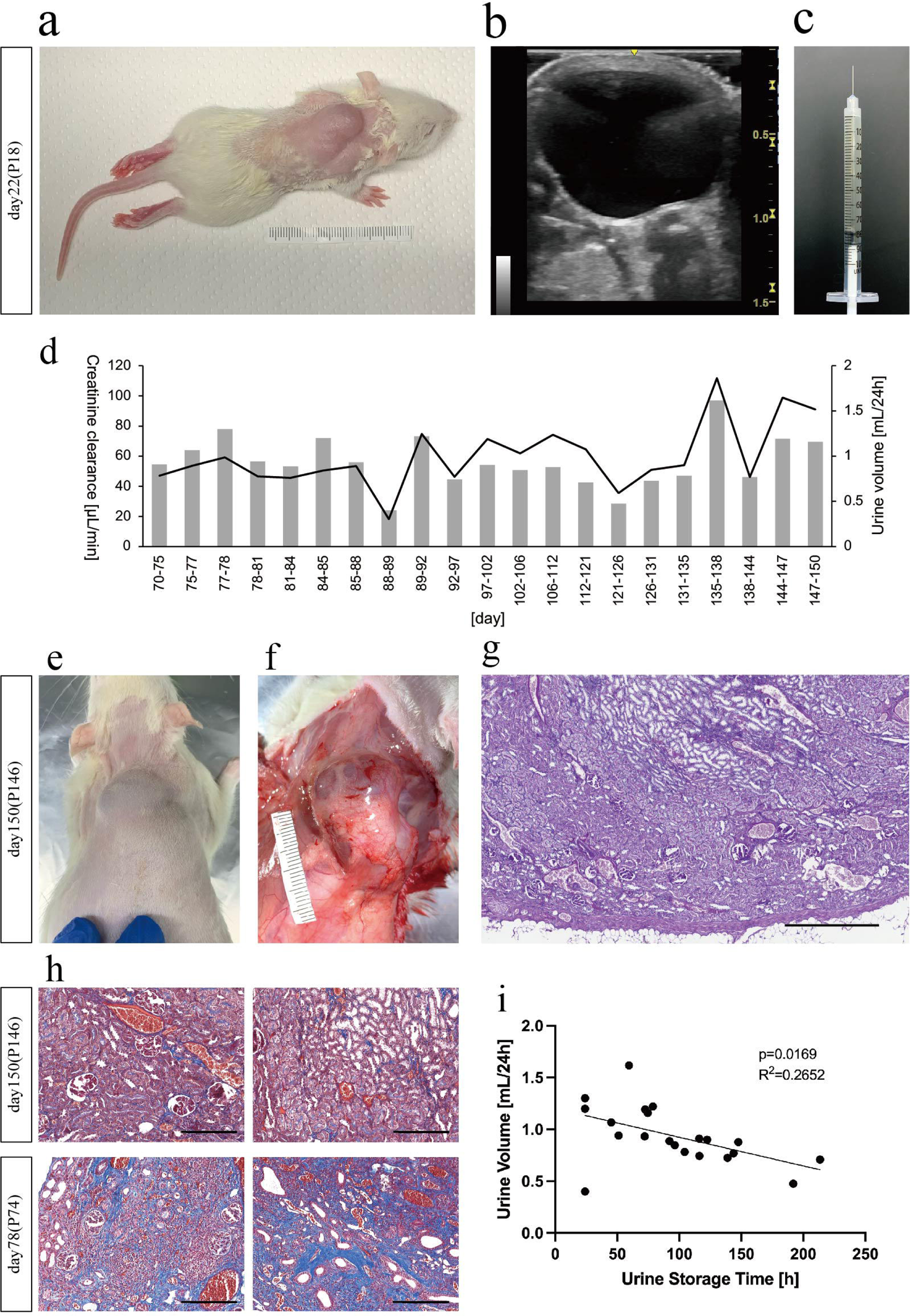
Development of the urine excretion method for GFP-SD rat MNBs transplanted into subcutaneous space. a At 22 days post-transplantation, a clearly raised structure, presumably the transplanted MNBs, was observed from the body surface. b Percutaneous ultrasonography revealed a distinct fluid accumulation. c A single puncture was performed from the back using a 29G needle, and urine was aspirated. d Bar graphs illustrating the volume of urine aspirated by puncture from transplanted MNBs, conducted at a frequency of 1-2 times per week, from day 70 to day 150 post-transplantation. Additionally, line graphs depicting the calculated creatinine clearance (n=1). e,f Tissue retrieval was performed at 150 days post-transplantation. g By periodic acid-Schiff staining, mature renal tissue structures including glomeruli and more developed tubules were confirmed. h Masson’s trichrome staining revealed a lack of fibrotic areas at 150 days post-transplantation, compared to tissue retrieval at 78 days post-transplantation without any aspiration punctures. i We re-plotted the data presented in Fig.3d as a scatter plot. The data were analyzed using the linear regression. A p-value of <0.05 was considered statistically significant. Scale bars, 500 μm in (g); 200μm in (h).

Compared to the tissue image at 28 days post-transplantation (Fig.2g), a more extensive mature tubular structure was observed (Fig.3g). Furthermore, Masson’s trichrome staining revealed a lack of fibrotic areas compared to tissue retrieval at 78 days post-transplantation without any aspiration punctures (Fig.3h).

Therefore, it was confirmed that repeated aspiration punctures for urine excretion prevented hydronephrosis and allowed for longer-term maturation of the MNBs. To investigate whether the frequency of 1-2 punctures per week was appropriate for optimal urine production, the correlation between the urine accumulation time between punctures (in this study, the minimum accumulation time between punctures was 24 hours) and the amount of urine produced per 24 hours was examined. It was found that shorter accumulation times tended to result in higher urine output per 24 hours (Fig.3i). On the other hand, to ensure reproducibility, similar MNBs transplantation experiments into fetuses were performed 17 times. However, in allogeneic transplantation, only 23.5% (4/17) of the individuals obtained a urinary cyst of sufficient size for aspiration puncture, as described above. In all experiments, tissue retrieval was performed more than 3 weeks after transplantation, and it was inferred that rejection occurred to varying degrees in individuals with poor urine production. This suggests that there are varying degrees of immunological advantage acquisition in fetal transplantation.

### 2.4 Verification of immunological advantages in fetal transplantation

Using fetuses with developing immune systems as recipients may have immunological advantages. To investigate whether tolerance is induced as one of the mechanisms, the following experiment was conducted. Fetal kidneys were individually extracted from GFP-SD rats (E14.0-14.5). One fetal kidney was transplanted into an SD rat fetus in utero (E18.0-18.5). The other fetal kidney from the same individual was then cryopreserved using a previously reported protocol 23). Four days after transplantation, on embryonic day 22, natural delivery occurred, and the pups were nursed by their mothers. One month later, they were weaned, and at 8 and 9 weeks after birth, when the immune system was fully mature, the cryopreserved fetal kidney was thawed and retransplanted into the retroperitoneal cavity of each individual. Tissue retrieval was performed 3 and 2 weeks after each transplantation (i.e., 11 weeks after birth) (Fig.4a). In the tissue retrieved 3 weeks after transplantation at 8 weeks of age, GFP-positive areas were sufficiently preserved (Fig.4b), but histologically, severe infiltration of inflammatory cells, mainly lymphocytes, was observed, and glomerular and tubular structures were barely visible due to rejection (Fig.4c,d,e). In contrast, in the tissue retrieved 2 weeks after transplantation at 9 weeks of age, GFP-positive areas were sufficiently preserved, and recipient-derived blood vessels were observed invading the fetal kidneys (Fig.4f,g). Histologically, although lymphocyte infiltration was observed, more glomerular and tubular structures were confirmed compared to the 3-week retrieval (Fig.4h,i,j). Compared to the rejection observed in the tissue retrieved 2 weeks after transplantation of GFP-SD rat MNBs into adult SD rats at 8 weeks of age, where glomerular and tubular structures could not be identified (Fig.2k), the rejection was clearly reduced.

**Figure 4.**
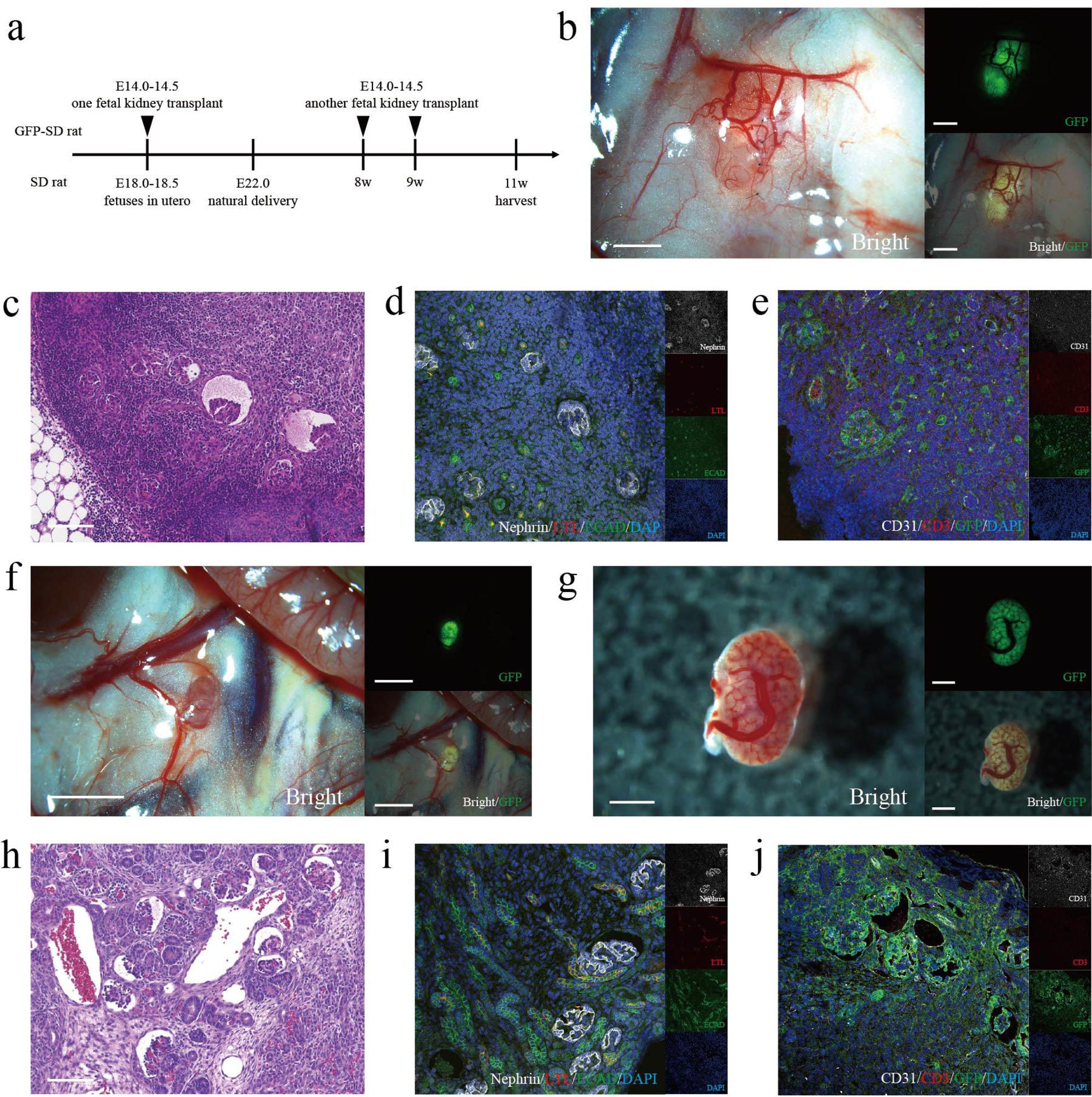
Verification of immunological advantages in fetal transplantation. a We illustrate the schematic diagram of the experiment in Fig.4a. When transplanting one fetal kidney (E14.0-14.5), the other fetal kidney is kept frozen. Eight and nine weeks after transplantation, the frozen kidney is thawed, re-transplanted, and tissues are retrieved 11 weeks after the first transplantation. b In the tissue retrieved 3 weeks after transplantation at 8 weeks of age, GFP-positive areas were sufficiently preserved. c,d,e By periodic acid-Schiff staining and immunofluorescence, severe infiltration of inflammatory cells, mainly lymphocytes, was observed, and glomerular and tubular structures were barely visible due to rejection. f,g In the tissue retrieved 2 weeks after transplantation at 9 weeks of age, GFP-positive areas were sufficiently preserved, and recipient-derived blood vessels were observed invading the fetal kidneys. h,i,j By periodic acid-Schiff staining and immunofluorescence, lymphocyte infiltration was observed, more glomerular and tubular structures were confirmed compared to the 3-week retrieval. Scale bars, 2 mm in (b) and (f); 1 mm in (g); 100 μm in (c) and (h).

### 2.5 Verification of xenotransplantation (mice to rats)

A similar experiment was conducted using xenotransplantation of GFP-B6 (C57BL/6) mouse fetal kidneys into SD rat fetuses. Four days after transplantation, on embryonic day 22, natural delivery occurred, and the pups were nursed by their mothers and grew. Tissue retrieval was performed after transplantation. Even without immunosuppressant administration, in the transplantation into SD rat fetuses, GFP-positive areas persisted 10 days after transplantation, and numerous CD3-positive cells infiltrated the tubular interstitial region, but glomerular structures were observed (Fig.5a,b). In contrast, when GFP-B6 mouse fetal kidneys were transplanted into adult SD rats at 8 weeks of age and tissue was retrieved 3 days after transplantation, severe rejection was observed before maturation (Fig.5c), suggesting that rejection is somewhat reduced even in xenotransplantation into fetuses. However, 18 days after transplantation, GFP-positive areas decreased, severe inflammatory cell infiltration was observed, and no glomerular structures were detected (Fig.5d,e). To reduce rejection, tacrolimus administration was performed. To transfer tacrolimus to the fetuses via the placenta, tacrolimus was subcutaneously administered to the mother rats at 0.2 mg/kg/day (every other day) after transplantation, and after birth, the neonates themselves were subcutaneously administered tacrolimus at 0.1 mg/kg/day (every other day). In a preliminary verification, the placental transfer of tacrolimus from the mother rat to the fetuses was evaluated, and the transfer rate to the fetuses was approximately 40%. With tacrolimus administration, GFP-positive areas were sufficiently preserved even 18 days after transplantation (Fig.5f), and histologically, glomerular and partial tubular structures were observed (Fig.5g). Immunostaining revealed ECAD-positive distal tubules but no LTL-positive proximal tubules, suggesting that xenotransplantation may affect tubular maturation (Fig.5h). The CD31-positive vessels inside the GFP-positive glomerular structures were GFP-negative, indicating that the glomeruli were completely composed of recipient (mouse)-derived vessels (Fig.5i). On the other hand, marked infiltration of CD3-positive cells was observed in the tubular interstitial region, confirming that tacrolimus monotherapy was insufficient to suppress rejection in xenotransplantation (Fig.5i).

**Figure 5.**
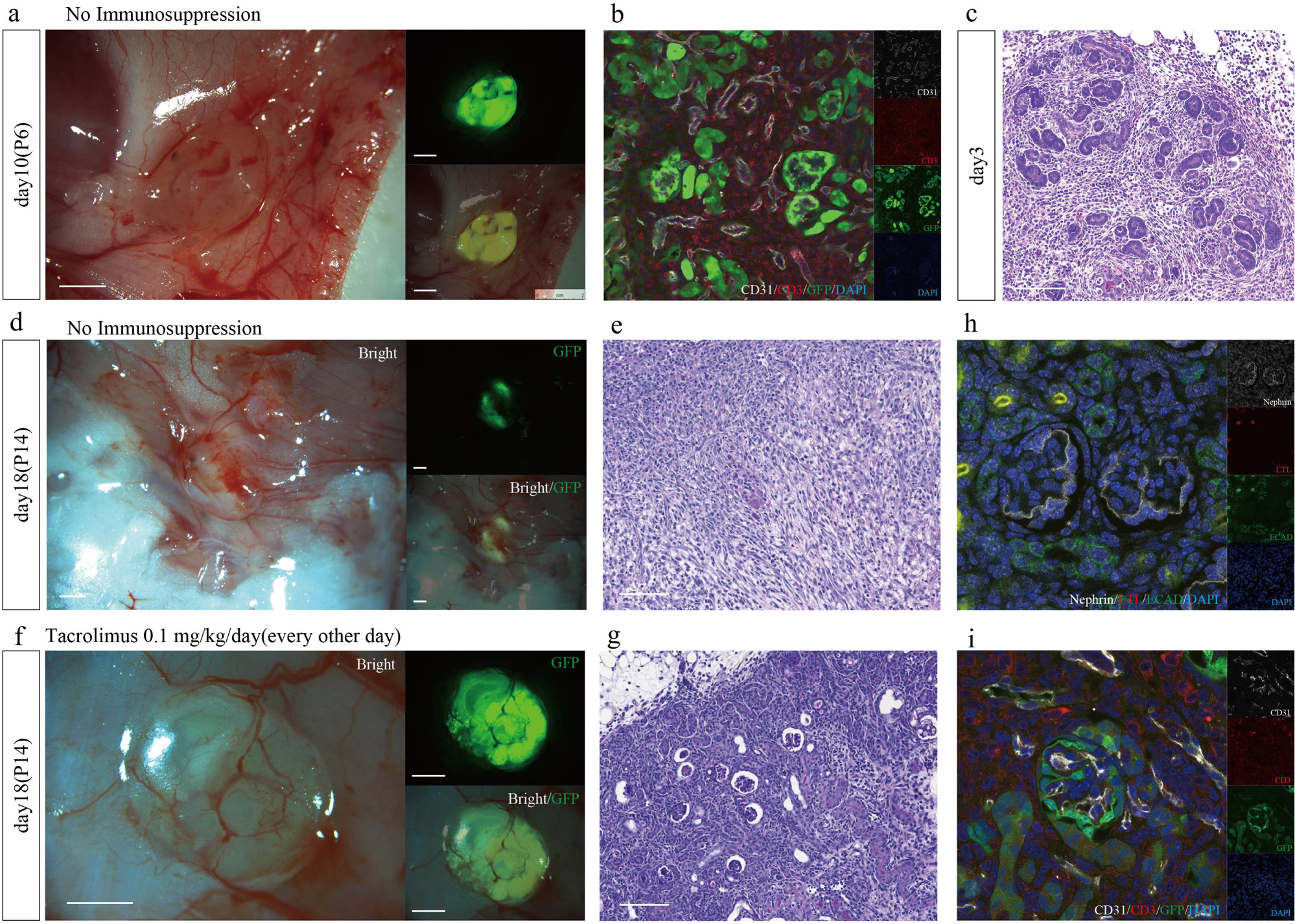
Verification of xenotransplantation (mice to rats) a,b No immunosuppression. Even without immunosuppressant administration, in the transplantation into SD rat fetuses, GFP-positive areas persisted 10 days after transplantation, and numerous CD3-positive cells infiltrated the tubular interstitial region, but glomerular structures were observed. c No immunosuppression. By hematoxylin and eosin staining, when a GFP-B6 mouse fetal kidneys was transplanted into adult SD rats at 8 weeks of age and tissue was retrieved 3 days after transplantation, severe rejection was observed before maturation. d,e No immunosuppression. Eighteen days after transplantation, GFP-positive areas decreased, severe inflammatory cell infiltration was observed, and no glomerular structures were detected (e periodic acid-Schiff staining). f With tacrolimus administration, GFP-positive areas were sufficiently preserved even 18 days after transplantation. g By periodic acid-Schiff staining, glomerular and partial tubular structures were observed. h Immunostaining revealed ECAD-positive distal tubules but no LTL-positive proximal tubules. i The CD31-positive vessels inside the GFP-positive glomerular structures were GFP-negative, indicating that the glomeruli were completely composed of recipient (mouse)-derived vessels. Marked infiltration of CD3-positive cells was observed in the tubular interstitial region. GFP, green fluorescent protein; LTL, Lotus tetragonolobus lectin; ECAD, E-cadherin. Scale bars, 1 mm in (a), (d), and (f); 100 μm in (c), (e), and (g).

## 3. Discussion

In this study, we successfully transplanted a urological unit: MNBs consisting of kidney, ureter and bladder into rat fetuses in utero using a needle via a transuterine approach. We obtained live offspring through natural delivery and created a mature exogenous kidney.

Previously reported transplantation techniques into the amniotic fluid 19) peritoneal cavity 20,21), or retroperitoneal space 22) of fetuses in utero involved the transplantation and administration of cells or liquids. In both cases, the invasiveness was minimized because thin puncture needles could be used. In contrast, we attempted organ transplantation as part of fetal therapy for the purpose of renal replacement therapy. For organ transplantation, a needle with a thickness capable of loading an organ inside was required. Therefore, as a measure to perform transplantation with a stable survival rate in rat fetuses, we developed a needle with the thinnest possible outer diameter and a thin wall thickness that could load an organ. We also focused on subcutaneous transplantation as the transplantation site. In previous studies using adult recipients in rodents, the retroperitoneal space was generally selected as the transplantation site. The retroperitoneal space can easily form a space for transplanting fetal organs by partially surgically peeling off the visceral peritoneum. However, transplantation into the retroperitoneal space requires laparotomy, making it unrealistic to perform transuterine transplantation. Interestingly, we discovered that subcutaneous transplantation 24), which was considered disadvantageous in adult transplantation, was suitable as a transplantation site for fetal organs in fetal transplantation. We have previously attempted to transplant fetal kidneys into the subcutaneous space of adults 24), but in all cases, the development was poor, and no significant urine production was obtained. We speculated that the reasons for the developmental insufficiency might be the strong immune barrier and the tight and narrow transplantation bed in the adult subcutaneous space, despite the presence of blood flow. In contrast, the subcutaneous space during the fetal period has a wider space to physically accommodate fetal organs compared to the retroperitoneal space, making it safer and easier to perform transuterine transplantation using a needle. Subcutaneous transplantation had a low incidence of bleeding complications, and the fetal survival rate was high at 76%. The expansion of the transplantation bed as the fetus grows may also alleviate physical constraints. In addition to the physical and spatial advantages, we hypothesized that there might be specific advantages of the fetal subcutaneous space, as there was a remarkable difference in the development of the transplanted MNBs after transplantation, with urine production, compared to adult recipient subcutaneous transplantation.

The fact that MNBs grow in the subcutaneous space of fetuses but not in the subcutaneous space of adults highlights the difference in the transplantation bed. For example, it has been reported that the extracellular matrix (ECM) composition differs between the subcutaneous space of adults and fetuses 25). The ECM of fetal skin has a higher ratio of type III collagen and glycosaminoglycans compared to that of adults. This difference is considered to be the reason for the different repair mechanisms between fetuses and adults, with fetuses having the ability to heal without scarring, and the involvement of ECM has been suggested as the cause 26). ECM is known to promote development and angiogenesis in kidney organoids that mimic embryonic organs 27). Additionally, some components of ECM have been reported to play a role in directing the migration of endothelial cells, contributing to angiogenesis 28). Thus, an analysis focusing on the unique ECM composition of the fetal subcutaneous space is necessary in the future to understand the difference between adult and fetal subcutaneous transplantation.

The production of such a large amount of urine for such a long period by a single MNB has not been observed in retroperitoneal transplantation in adults. Not only were fetuses suitable as recipients for fetal organ transplantation, but we also consider that the ease of urine excretion management through subcutaneous transplantation was an important advantage, as regular urine excretion is crucial for preventing hydronephrosis in fetal kidney transplantation.

Once urine is produced from the fetal kidneys, if left untreated, the transplanted MNBs will develop hydronephrosis due to the lack of a urine excretion pathway, and eventually, renal function will be lost. Therefore, the formation of a urine excretion pathway becomes necessary at 3-4 weeks after transplantation when urine production is observed. In previous studies, MNBs were transplanted into the retroperitoneal space of adults, making it difficult to excrete urine through percutaneous aspiration from the body surface. Therefore, a surgical method was employed to connect the urinary cyst to the host’s ureter by laparotomy 8). In our model, we took advantage of the fact that the transplanted MNBs were located subcutaneously and in close proximity to the body surface. We successfully performed safer and continuous percutaneous aspiration for a long period of 150 days, avoiding hydronephrosis and achieving urine excretion to the outside of the body.

In this study, we performed aspiration punctures 1-2 times per week because we determined that a certain amount of urine needed to accumulate for safe aspiration. After aspirating the urine, it re-accumulates in the bladder over a certain period, but the optimal frequency of aspiration punctures for better urine production was unknown. Therefore, we investigated the appropriate frequency of aspiration punctures and found that shorter accumulation times between punctures tended to result in a larger amount of urine produced. This suggests that more continuous urine excretion, rather than intermittent excretion, may result in a higher urine output per unit time. Considering safety, we performed aspiration punctures 1-2 times per week in this study, but in the future, we aim to establish a method for continuous drainage, such as by placing a catheter.

Moreover, when considering long-term engraftment, it is important to consider immunological advantages.

In the allogeneic transplantation of MNBs into fetuses (SD rat-SD rat), the rejection was reduced to such an extent that urine production persisted for 150 days without the use of immunosuppressants. This may be attributed to two advantages of fetal kidney transplantation into fetuses in utero. One is that the donor organ was fetal kidneys, as previously reported 29,30), and the other is that the recipient was a fetus, which is a characteristic of this study. Regarding the former, it has been reported that fetal kidneys may have lower immunogenicity than adult-type kidneys 29,30). Furthermore, fetal kidneys are avascular at the time of transplantation, but as they grow after transplantation, recipient-derived blood vessels invade and form glomeruli. In other words, the fact that the donor organ’s vasculature is constructed by the recipient side is considered an advantage, as the blood vessels are less likely to be targeted by rejection 17). In this study, the vasculature of the transplanted fetal kidneys was observed as recipient-derived vessels expressing CD31 without merging with GFP. On the other hand, partial donor-derived vessels expressing GFP were also observed. It was speculated that vascular endothelial progenitor cells 31) present in the fetal kidneys differentiated and partially constructed chimerized vessels 32,33). At least, since fetal kidneys are composed of recipient-derived vessels, it is conceivable that rejection is reduced compared to organ transplantation using adult organs as donors.

Another immunological advantage is that the recipient is a fetus, which is a characteristic of our approach. Transplantation during the fetal period, when the immune system is still developing, may have immunological advantages. Fetuses develop immune tolerance to self-antigens and maternal antigens while growing in the maternal uterus 34). Attempts have been made to induce tolerance to foreign antigens by transplanting them during the fetal period, but the degree of tolerance acquisition varies, and consistent results have not been obtained 35,36,37). In the allogeneic transplantation in this study, the difference in the degree of tolerance may have influenced the difference in the degree of urine production. However, good histological findings were observed, suggesting that transplantation at an early stage of fetal development may have induced partial tolerance and reduced the rejection response. On the other hand, in xenogeneic transplantation into fetuses, long-term engraftment could not be achieved without the use of immunosuppressants. However, when comparing adult and fetal recipients, in the xenogeneic fetal kidney transplantation, severe rejection was confirmed at day 3 in adults, while glomeruli were still observed at day 10 in fetal transplantation. Furthermore, the addition of a small amount of immunosuppressant resulted in longer-term engraftment (18 days). Previous reports have shown that cell transplantation can engraft in fetuses in utero in xenogeneic settings 38,39,40). The immunological immaturity of the recipient may be advantageous for xenotransplantation. Although xenotransplantation has a higher hurdle in terms of rejection compared to allogeneic transplantation, immunological advantages through the above-mentioned mechanisms were speculated.

A limitation of this study is that a kidney failure model was not used as the transplantation target. Because all recipient fetuses were normal (wild type) fetuses, the actual therapeutic effect of fetal kidney transplantation could not be confirmed. In the future, we plan to perform transplantation verification using a congenital kidney disease model (Six2-expressing nephron progenitor cell depletion model) in rats. Furthermore, the CCr of a single MNBs is still low at 40-80 μL/min compared to the CCr of 4000 μL/min calculated from the 24-hour urine collection of the neonates themselves at the same time point. To achieve therapeutic effects in the future, it is necessary to explore methods to enhance the function of a single MNBs or to transplant multiple MNBs. Additionally, it is necessary to determine the minimum number of MNBs and the transplantation period required for bridging to dialysis. Regarding the timing of transplantation in recipients, performing transplantation at an earlier stage may provide immunological tolerance advantages. Although there are size limitations in rodents, transplantation at an earlier developmental stage may be possible in larger animals or humans. To aim for clinical application, technical investigations are also being conducted in pig fetuses as large animals. As the next stage, immunological analyses will be performed using non-human primates as recipients.

The ultimate goal of this study is to develop renal replacement therapy for children with severe congenital kidney disease, like Potter sequence, for which there are few treatment options. In this study, we used MNBs as transplantation organs, established a transplantation method into fetuses, and demonstrated for the first time in the world the success of transplantation into fetuses and the acquisition of functional organs capable of urine production in allogeneic settings. The superiority of fetal tissue as a donor organ has been previously pointed out, but this study suggests that the fetal elements on the recipient side may also have immunological and scaffold environment advantages. The results obtained in this study have the potential to be utilized in adult xenotransplantation research, which has made remarkable progress in recent years, application to other organs such as the heart and liver, and organoid research by analyzing the mechanisms from the perspective of fetal specificity. In this study, we succeeded in constructing an organ with urine production ability for as long as 150 days, which is considered a sufficient period as a bridge to dialysis in neonates, considering the lifespan of rodents. For application to humans, it is necessary to demonstrate the therapeutic effect using larger animal models, increase the number of transplanted MNBs, and improve the quality of transplanted MNBs. This study is an important step toward the development of a new treatment method for medical intervention in children with severe kidney disease, for whom no effective treatment has been available.

## 4. Methods

### 4.1 Research Animals

Animal experiments followed the Guidelines for the Proper Conduct of Animal Experiments of the Science Council of Japan (2006) and were approved by the Institutional Animal Care and Use Committee of the Jikei University School of Medicine (protocol numbers: 2023-021). All efforts were made to minimize animal suffering. Pregnant female Sprague-Dawley (SD) rats, Sprague-Dawley-Tg (CAG-EGFP [enhanced green fluorescent protein]) rats (GFP-SD rats), C57BL/6-Tg (CAG-EGFP) mice (GFP-B6 mice) and adult male SD rats were purchased from Sankyo Labo Service Corporation (Tokyo, Japan).

### 4.2 Isolation of fetal kidneys or MNBs

Pregnant rats and mice were anesthetized by isoflurane (2817774; Pfizer, New York, USA) inhalation. Fetal kidneys or MNBs (E14.0-16.5 of rats and E13.0-13.5 of mice) were harvested, and the pregnant rats and mice were then killed immediately by an infusion of pentobarbital (120 mg/kg). All the Fetuses were killed by decapitation. Fetal kidneys or MNBs were dissected under a surgical microscope (M205FA; Leica Microsystems, Wetzlar, Germany).

### 4.3 Transplantation of Rodent MNBs into the Fetal Subcutaneous Space

A summary of this experiment is shown in Fig.1a. GFP-SD rat MNBs (E14.0-16.5) (or GFP-B6 mouse fetal kidneys) were removed and loaded onto the tip of the 15 or 16G needle (Saito Medical Instruments Inc., Tokyo, Japan) (Fig.1b). The opposite end of the 15 or 16G needle was attached to a 0.5 mL syringe filled with HBSS (TERUMO, Tokyo, Japan). Pregnant SD rats (E18.0-18.5) were anesthetized by isoflurane inhalation. Subsequently, a midline incision was made, and the uterus was gently extracted from the abdominal cavity. The position of the fetuses was identified through the translucent uterine wall (Fig.1c). Using the 15 or 16G needle loaded with GFP-SD rat MNBs (or GFP-B6 mice fetal kidneys), the uterine wall was punctured on the dorsal side of the fetus, and while confirming under a stereomicroscope, the needle was inserted directly into the fetal subcutaneous space (Fig.1d,e). The needle was advanced approximately 5 mm under the fetal skin, and 0.1-0.5 mL was ejected to transplant the GFP-SD rat MNBs (or GFP-B6 mice fetal kidneys) into the fetal subcutaneous space. Immediately after transplantation, the needle was withdrawn. The presence of GFP-positive tissue was confirmed by fluorescence stereomicroscopy immediately after transplantation to ensure proper execution (Fig.1f). To minimize surgical duration, 2-4 fetuses per dam were used as recipients. The uterus was gently returned to the abdominal cavity. Approximately 5 mL of HBSS warmed to approximately 30°C was administered intraperitoneally to prevent adhesions. The anatomical positioning of the uterus within the abdominal cavity was confirmed, and the incision was closed.

### 4.4 Postnatal Management

Following natural birth at embryonic day 22, neonates were nursed by their biological mothers in the same cage. At postnatal day 0, the GFP-positive tissue was visualized on the neonates’ body surfaces using fluorescence microscopy. Neonates lacking the GFP-positive tissue was euthanized on the same day by decapitation. Body weight measurements were regularly conducted throughout the observation period.

Additionally, the GFP-positive tissue was visualized using fluorescence microscopy, and the longitudinal diameter was measured over time. When necessary, fetal kidneys or MNBs transplanted were observed using transcutaneous ultrasonography (LOGIQ e Premium; GE HealthCare Technologies Inc, Chicago, USA). If a urinary cyst with a longitudinal diameter of 5mm or more was detected via echography, it was deemed safe for percutaneous aspiration. Neonates were anesthetized by isoflurane inhalation, and a single puncture was made using a 23-29G needle (TERUMO, Tokyo, Japan) on the dorsal side to aspirate urine. Single punctures were repeated at a frequency of 1-2 times per week, and urine volume was measured.

### 4.5 Cryopreservation and Thawing Methods for fetal kidneys

The cryopreservation and thawing methods for fetal kidneys have been previously reported (Kenji Matsui et al. J Clin Med. 2023 Mar 15;12(6):2293. doi: 10.3390/jcm12062293. Cryopreservation of Fetal Porcine Kidneys for Xenogeneic Regenerative Medicine). Initially, the fetal kidneys were equilibrated in a base medium (MEM α supplemented with 20% fetal bovine serum [FBS; SH30070.03, HyClone Laboratories, Inc., Logan, UT, USA] and 1% antibiotic–antimycotic solution [15,240,062; Thermo Fisher Scientific, Waltham, MA, USA]) containing 7.5% ethylene glycol (EG; 055-00996; Wako, Osaka, Japan) and 7.5% dimethyl sulfoxide (DMSO; 317275-100ML; Millipore, Burlington, MA, USA) on ice for 15 minutes, followed by soaking in a base medium with 15% EG and 15% DMSO on ice for an additional 15 minutes. Subsequently, the fetal kidneys were placed onto Cryotops (81111; Kitazato Corporation, Tokyo, Japan) and immediately submerged into liquid nitrogen for storage until transplantation. Fetal kidneys cryopreserved for several weeks were thawed just prior to transplantation. The Cryotops containing fetal kidneys were swiftly transferred from liquid nitrogen to a base medium with 1 M sucrose at 42°C for 1 minute, then to a base medium with 0.5 M sucrose at room temperature for 3 minutes, and finally washed twice in a base medium at room temperature for 5 minutes each time.

### 4.6 Transplantation of fetal kidneys or MNBs

SD rats were anesthetized by isoflurane inhalation, and a laparotomy was performed through an abdominal midline incision. A pocket was created in the retroperitoneal space within the region bounded by the aorta, left ureter, and left renal artery, using micro-tweezers (11253-25; Dumont, Montignez, Switzerland) under a surgical microscope. A single fetal kidneys or MNB was transplanted into the pocket, which was subsequently closed using 10-0 nylon sutures (Muranaka Medical Instruments Co. Ltd., Osaka, Japan). The procedure was completed by closing the abdominal incision.

### 4.7 Biochemical Measurements of Blood and Urine

For blood tests, SD rats were anesthetized by isoflurane inhalation, and their tail veins were punctured using a 25G needle (TERUMO) to collect blood samples using a hematocrit capillary tubes (2-454-21, AS ONE Corporation). The capillaries were sealed with wax (2-454-22, AS ONE Corporation) and centrifuged at 12,000 g for 10 minutes to separate serum and blood cells. Serum creatinine levels were quantified using a DRI-CHEM (FUJIFILM). Urine samples were collected from MNBs and from the rats’ 24-hour urine collection. For the 24-hour urine collection, SD rats were placed in rat metabolic cages (KN-650-MC, Natsume Seisakusho Co., Ltd.) overnight. The urine samples were submitted to SRL, Inc. for measurement of urine creatinine levels.

### 4.8 Histology and Immunofluorescence

The harvested fetal kidneys or MNBs were fixed in 4% paraformaldehyde (161-20141, Wako) at 4°C overnight and subsequently embedded in paraffin. Sections of 4 μm thickness were obtained using a rotary microtome (HM355S, PHC, Tokyo, Japan). Standard procedures were followed for staining with hematoxylin and eosin (HE), Masson’s trichrome, and periodic acid-Schiff (PAS).

For immunostaining, the slides were deparaffinized, washed thrice in phosphate-buffered saline (PBS), and subjected to antigen retrieval by incubating with citrate buffer (#K 035; 10X Citrate Buffer, pH 6.0, DBS) at 121°C for 10 minutes. Following three additional washes in PBS, the slides were blocked at room temperature for 30 minutes using 5.0% skimmed milk. Subsequently, the sections were incubated overnight at 4°C with primary antibodies (Table.2), followed by incubation with secondary antibodies conjugated with Alexa Fluor 488, 546, or 647, and DAPI for 1 hour at room temperature. Finally, the sections were mounted with a glycerol-based liquid mounting medium. Each sample was examined under a fluorescence microscope (LSM880; Carl Zeiss, Oberkochen, Germany).

**Table 2.** List of primary antibodies.

| **Antigen** | **Host** | **Supplier** | **Cat. No.** | **Dilution** |
| --- | --- | --- | --- | --- |
| Nephrin | Guinea pig | Progen | GP-N2 | 1:100 |
| LTL biotin | - | Vector | B-1325 | 1:200 |
| E-cadherin | Mouse | Thermo | 610181 | 1:100 |
| E-cadherin | Rabbit | Cellsignaling | 3195S | 1:100 |
| GFP | Rabbit | MBL | MBL598 | 1:500 |
| GFP | Chicken | Abcam | ab13970 | 1:100 |
| CD31 | Goat | R&D | AF3628 | 1:100 |
| CD3 | Mouse | DAKO | Clone F7.2.38 | 1:100 |
| CD3 | Rabbit | Abcam | ab5690 | 1:100 |

### 4.9 Statistical Analyses

All data are presented as mean ± standard error of the mean (SEM). The data were analyzed using the two-tailed unpaired t-test and linear regression. A p-value of <0.05 was considered statistically significant. Experimental data were analyzed using GraphPad Prism version 8.0 (GraphPad Software, Boston, MA, USA) and Microsoft Excel (Microsoft, Redmond, WA, USA). Details are explained in each figure legend.

## Acknowledgments

We thank Ms. M. Tamatsukuri, Ms. T. Hayakawa, Ms. S. Kawagoe, and Ms. H. Gotoh for their experimental and technical assistance. This work was supported by the Japan Agency for Medical Research and Development (AMED; grant no. 22bm0704049h0003 and 24bm1223003h0003).

## Author contributions

K.Morimoto. wrote the main manuscript text. K.Morimoto and S.Yamanaka designed the study. K.Morimoto performed the experiments. K.Morimoto, S.Yamanaka, S.Yamamoto, N.K., Y.K., Y.I., K.Matsui, N.M., Y.S., T.T., T.F., S.F., S.T., K.Matsumoto, K.O., S.W., E.K., T.Y. interpreted the data and revised the manuscript. S.Yamanaka, E.K. and T.Y. supervised the study. All authors reviewed the manuscript and granted permission to publish the study. T.Y. granted final permission to the manuscript.

## Data availability statement

The main data supporting the results in this study are available within the paper and its Supplementary Information. All other data are available from the corresponding author upon reasonable requests.

## Additional Information

Correspondence and requests for materials should be addressed to Takashi Yokoo. Conflicts of Interest: Author Eiji Kobayashi is the director of Kobayashi Regenerative Research Institute, LLC. Eiji Kobayashi has received a research fund from Sumitomo Pharma Co., Ltd., Osaka, Japan as a result of the Collaborative Research Agreement between The Jikei University School of Medicine and Sumitomo Pharma Co., Ltd. Author Eiji Kobayashi have received technical guidance fee from Fuji Micra Inc., Shizuoka, Japan. The authors declare that this research was conducted in the absence of any commercial or financial relationships that could be construed as a potential conflict of interest. The funders had no role in the design of the study; in the collection, analyses, or interpretation of data; in the writing of the manuscript, or in the decision to publish the results.

